# A Critical Appraisal of Predatory Journals in Pathology

**DOI:** 10.1101/482174

**Authors:** Yaman M. AlAhmad, Ibrahim Abdelhafez, Farhan S. Cyprian, Faruk Skenderi, Saghir Akhtar, Semir Vranic

**Author notes:** Correspondence: Semir Vranic, MD, PhD, College of Medicine, Qatar University, 2713 Doha, Qatar, or. These authors contributed equally (Yaman M. AlAhmad, Ibrahim Abdelhafez).

## Abstract

Predatory or pseudo journals have recently come into focus due to their massive internet expansion and extensive spam email soliciting. Recent studies explored this urging problem in several biomedical disciplines. In the present study, we identified 69 potential predatory (pseudo) pathology journals that were contrasted to 89 legitimate pathology journals obtained from the major bibliographic databases. All potential predatory journals in pathology shared at least one of the features proposed by previous studies (e.g. a poor web-site integrity, submissions via email, unclear or ambiguous peer-review process, missing names of the editorial board members, missing or pending the journal ISSN). Twenty-one (30%) of the potential predatory pathology journals had misleading titles mimicking those of legitimate journals. Only one of the identified journals was listed in the Directory of Open Access journals whereas none (0%) was indexed in PubMed/MEDLINE or Web of Science, listed in the Committee on Publication Ethics nor have they had a legitimate impact factor in the Journal Citation Reports.

## Introduction

Predatory or pseudo journals refer to journals that recruit articles through aggressive marketing and spam emails, promising quick, but not robust review and fast open-access (OA) publication, thus compromising scholarly publishing standards (1-4). Their key motive is a financial benefit via article processing charges (APCs) and other additional fees (1, 3, 4).

The number of OA journals has dramatically risen over the past fifteen years (5), reaching 11,376 journals, indexed in the Directory of Open Access Journals (DOAJ) in 2018 (available at https://doaj.org). This expansion was parallel to the increase in the number of predatory publishers from 18 in 2011 to more than 1100 in 2016 (6, 7). Predatory (pseudo) journals have become more prevalent than ever due to massive internet expansion and extensive spam email soliciting (2, 4, 8). These journals are thought to exert a negative impact on the science including biomedicine.

Since 2011, when Jeffrey Beall, an academic librarian from the University of Colorado Denver, posted his first list of potential predatory, OA publishers and journals (available at: https://beallslist.weebly.com), predatory journals have come into focus (3, 4).

Recent studies have highlighted the significant impact of potentially predatory journals in several biomedical fields including neuroscience/neurology, urology, emergency medicine, physical medicine, orthopedics, anesthesiology and pediatrics (9-11). These studies revealed that by October 2016 >10% of predatory journals in three biomedical fields were indexed in PubMed (12% for rehabilitation, 11.4% for neurosciences, and 20.2% for neurology). By April 2017, these values increased to 23.7% for rehabilitation, 16.1% for neuroscience, and 24.7% for neurology, indicating that these journals evolved rapidly (6, 7). The study on emergency medicine revealed that almost half (25/55) of the OA journals were predatory journals (12). Furthermore, the study on urology found that (7/32) potential predatory journals, which were solicited to an academic urologist, were indexed in reputable databases including: Journal Citation Reports (JCR), Scimago Journal Rankings (SJR), and DOAJ (13). In contrast, a recent study of Kokol et al. exploring predatory journals in pediatrics revealed 26 such journals; however, none of them were indexed in PubMed, Scopus or Web of Science (9).

The role and presence of potential predatory journals in pathology have not been explored so far. In this study, these journals have been critically evaluated and were contrasted to the legitimate pathology journals.

## Materials and Methods

### 1. Journals Identification and Selection

Between January and May 2018, Beall’s list of predatory journals served as an initial database of suspected journals related to pathology, as it was previously used in other studies (2, 7). The term “potential predatory” was only used after assessing each journal separately based on the recommended criteria that are summarized in Table 1 (2, 8). On the other hand, the major bibliographic databases were explored (PubMed/MEDLINE, PubMed Central (PMC), and Web of Science/Science Citation Index (Science Citation Index/Science Citation Index Expanded (SCI/SCIE)) to identify legitimate journals in pathology. The journal titles were retrieved using the following key words: anatomic pathology, cellular pathology, clinical pathology, cytopathology, diagnostic pathology, experimental pathology, histopathology, human pathology, immunopathology, medical pathology, molecular pathology, molecular biomarkers, neuropathology, pathology, and surgical pathology.

**Table 1:** Features of potential predatory journal as proposed by Clemons et al. and Shamseer et al. (2, 8)

|  | Features |
| --- | --- |
| 1 | Set up journal names and websites that resemble or appear to be tied to those of legitimate ones |
| 2 | Wide scope of interest covering biomedical and non-biomedical subjects |
| 3 | Having spelling and grammar mistakes in the website |
| 4 | Displayed images are usually distorted and fuzzy |
| 5 | The website homepage language targets authors |
| 6 | Stating plans for indexing in reputable databases such as PubMed, while emphasizing being indexed in Index Copernicus. |
| 7 | Unreal number of issues per year, missing or pending ISSN number |
| 8 | Absent and/or unclear peer-review process |
| 9 | Manuscripts are submitted via email |
| 10 | Fast publication is assured |
| 11 | Absence of retraction and/or copyright policies |
| 12 | Lack of journal archive and/or digital preservation of content. |
| 13 | Low APCs (Article Processing Charges) |
| 14 | Personalized spam emails targeting a variety of authors inviting them to contribute with short deadlines |
| 15 | The contact email is non-journal affiliated (e.g., @hotmail.com) |

### 2. Data Collection (phase 1)

After collecting the available number of journals related to pathology, each journal was assessed based on, Clemson and colleagues’ criteria; its indexing status, clarity of peer-review process, availability of its archive, legitimacy of editorial board, the status of International Standard Serial Number (ISSN), emphasis on OA, and country of origin (8).

#### 2.1 Status of ISSN and country of origin

To identify the journal’s country of origin and the ISSN legitimacy of the suspected journals, the ISSN Portal service was used (available at https://portal.issn.org), which is an online database that provides information about the journal’s resource and record including the country of origin.

#### 2.2 Amount of APCs

In order to obtain the amount of requested APCs, each journal’s web site was checked for the Creative Commons ‘Attribution’ License Non-Commercial (CC BY-NC) price. Furthermore, APCs of other currencies were converted to the US Dollar (USD) using the currency converter (available at https://xe.com).

#### 2.3 Emphasis on OA, legitimacy of editorial board, and clarity of peer-review process

Journals that were promoting OA publishing modality on their websites’ homepage, displaying phrases similar to “Promoting OA, Supports OA, Your Way To OA”, were labeled under “emphasis on open-access”. Editorial boards were reviewed in each journal and a missing editorial board was labeled. Unclear peer-review process was labeled under “ambiguous or unclear peer-review process”.

#### 2.4 Website integrity, number of issues, and misleading titles

For the website integrity, marking was based on presence of major design flaws, compatibility and language mistakes. “Unreal or small number of issues per year” was determined by comparing displayed number with the available number. To identify misleading journal names, a minimum of two identical words was required.

#### 2.5 Indexing status

The indexing status displayed on the journal’s website was appraised via both the National Library of Medicine’s (NLM) catalog (available at https://ncbi.nlm.nih.gov/nlmcatalog/) and the DOAJ.

### 3. Data Collection (phase 2)

After obtaining preliminary data from phase 1 of the assessment, each journal was further investigated based on Shamseer and colleagues’ criteria; aims and scope, journal name and publisher, home page integrity, contact information, indexing and impact factor, editorial processing and peer review, publication ethics and policies, and publication model and copyright (2). Since DOAJ indexing status, ISSN, APCs and misleading titles were obtained in phase 1, their findings were only reviewed.

#### 3.1 Aims and scope, home page integrity, and contact information

The aims and scope were investigated for inclusion of non-biomedical subjects. Spelling and/or grammar errors, distorted images, author-targeting homepage language, and non-journal affiliated emails, were all collected.

#### 3.2 Editorial processing, and publication copyright, ethics and policies

Presence of editor-in-chief, editorial board, submission system, peer review, retraction, copyright and plagiarism policies, and APC amount, were labeled accordingly.

#### 3.3 Indexing and impact factor

The status of each journal was checked in PubMed/Medline, Web of Science (SCI/SCIE), the World Association of Medical Editors (WAME) (available at http://wame.org), and the Committee on Publication Ethics (COPE) (available at https://publicationethics.org/members). Additionally, the Index Copernicus value was also considered.

### 4. Statistical Analysis

Shapiro–Wilk test and histogram were used to check for normality. The non-parametric, Mann-Whitney test was used to determine statistical significance of the APC means between OA legitimate and potential predatory journals. All analyses were performed using SPSS Statistics 25.0 for Windows (IBM, Armonk, NY, USA). Statistical significance was accepted at *P* < 0.05.

## Results

Of the total current publishers on Beall’s list (1196), 69 potential predatory journals were identified in the field of pathology (“the black list” journals). The journals’ titles and their respective publishers are shown in Table 2. Only one of the identified potential predatory journals in pathology (*Journal of Modern Human Pathology*) was indexed in the DOAJ. None of these journals were indexed in PubMed/MEDLINE, Web of Science (SCI/SCIE) nor listed in COPE or had a legitimate impact factor in the JCR (Clarivate Analytics) (“the white list” journals). Of note, 13 potential predatory journals in pathology had at minimum one of their articles archived in the PMC repository (following the policy of PMC as a digital repository archiving free full-text articles that had been published within the biomedical and life science journals). On the other hand, 89 legitimate journals were identified in the field of pathology and used for the comparison (listed in Table 3).

**Table 2:** A list of potential predatory journals in pathology (n = 69)

|  | Journal | Publisher |
| --- | --- | --- |
| 1 | Academic Open Clinical Pathology Research Journal | Academic Knowledge and Research Publishing |
| 2 | Advanced Journal of Surgical Pathology | Advanced Scholars Journals |
| 3 | African Journal of Cellular Pathology | Clavenanum Press Nigeria Limited |
| 4 | Allied Journal of Clinical Pathology Research | Allied Academies |
| 5 | American Research Journal of Pathology | American Research Journals |
| 6 | Annals of Clinical Pathology Research | Remedy Publications |
| 7 | Annals of Clinical Pathology | JSciMed Central |
| 8 | Archives of Pathology and Clinical Research | Heighten Science Publications |
| 9 | Archives of Clinical Pathology | SciTechnol |
| 10 | Asian American Pathology Research Journal | Asian and American Research Publishing Group |
| 11 | Austin journal of clinical pathology | Austin Publishing Group |
| 12 | Austin Journal of Pathology & Laboratory Medicine | Austin Publishing Group |
| 13 | Journal of Clinical Pathology and Forensic Medicine | Academic Journals |
| 14 | BAOJ Pathology and Clinical Research | Bio Accent |
| 15 | British Open Journal of Pathology | British Open Research Publications |
| 16 | Canadian Open Clinical Pathology Journal | Canadian Research Publication |
| 17 | Case Reports in Clinical Pathology | Sciedu Press |
| 18 | Journal of Clinical and Diagnostic Pathology | Open Access Pub |
| 19 | International Journal of Clinical and Experimental Pathology | e-Century Publishing Corporation |
| 20 | Clinical and Diagnostic Pathology | Open Access Text (OAT, OA Text) |
| 21 | Journal of Clinical & Experimental Pathology | OMICS International |
| 22 | CRESSCO International Journal of Pathology | Cresco Online Publishing |
| 23 | Current Updates in Clinical Pathology | OPR Science |
| 24 | Developments in Clinical & Medical Pathology | Crimson Publishers |
| 25 | Diagnostic Pathology: Open Access | OMICS International |
| 26 | Donnish Journal of Clinical Pathology and Forensic Medicine | Donnish Journals |
| 27 | Eurasian Clinical Pathology Research Journal | Eurasian Research Publishing |
| 28 | European Open Clinical Pathology Journal | European Union Research Publishing |
| 29 | Global Scientific Research Journal of Pathology | Global Scientific Research Journals (GSR) |
| 30 | HSOA Journal of Pathology Clinical & Medical Research | Herald Scholarly Open Access |
| 31 | Immunochemistry & immunopathology | OMICS International |
| 32 | Integrative Clinical Pathology | Scient Open Access |
| 33 | International clinical pathology journal | MedCrave |
| 34 | International Journal of Clinical Pathology and Diagnosis | Gavin Publishers |
| 35 | International Journal of Clinical Pathology Research | Academic and Scientific Publishing |
| 36 | International Journal of Microbiology and Pathology | OMICS International |
| 37 | Journal of Speech Pathology & Therapy | OMICS International |
| 38 | International Journal of Pathology and Clinical Research | ClinMed International Library |
| 39 | International Journal of Pathology Research and Practice | Axis Journals |
| 40 | IP Journal of Diagnostic Pathology and Oncology | Innovative Publication |
| 41 | The open forensic science journal | Bentham Open |
| 42 | Journal of Cellular & Molecular Pathology | Insight Medical Publishing (IMedPub) |
| 43 | Journal of Clinical & Anatomic Pathology | JScholar Journals |
| 44 | Journal of Clinical & Experimental Pathology | OMICS International |
| 45 | Journal of Clinical Pathology and Cytology | Bio Accent |
| 46 | Journal of Immunopathology | Pulsus Group |
| 47 | Journal of Medical & Surgical Pathology | OMICS International |
| 48 | Journal of Medicine, Radiology, Pathology and Surgery | Incessant Nature Science Publishers (INSPublishers) |
| 49 | Journal of modern human pathology | NobleResearch |
| 50 | Journal of Molecular Biomarkers & Diagnosis | OMICS International |
| 51 | Journal of MPE Molecular Pathological Epidemiology | Insight Medical Publishing (IMedPub) |
| 52 | Journal of Pathology and Therapeutics | Sci Forschen |
| 53 | Pathology and Laboratory Medicine | Science Publishing Group |
| 54 | North American Open Clinical Pathology Research Journal | North American Research Publishing |
| 55 | North American Open Physiology & Pathophysiology Research Journal | North American Research Publishing |
| 56 | Open Journal of Pathology | Scientific Research Publishing (SCIRP) |
| 57 | Pathology and Laboratory Medicine - Open Journal (PLMOJ) | Openventio Publishers |
| 58 | Pathology Discovery | Herbert Open Access Journals |
| 59 | International Journal of Ophthalmic Pathology | SciTechnol |
| 60 | Recent Advances in Pathology & Laboratory Medicine (RAPL) | Advanced Research Publications |
| 61 | Reports in Disease Markers | OMICS International |
| 62 | Saudi Journal of Pathology and Microbiology | Scholars Middle East Publishers |
| 63 | Swift Journal of Clinical Pathology and Forensic Medicine | Swift Journals |
| 64 | The Internet Journal of Pathology | Internet Scientific Publications |
| 65 | TJPRC: Journal of Human Pathology & | Trans Stellar |
|  | Research |  |
| 66 | Universal Open Forensic Medicine and Pathology Journal | Adyan Academic Press |
| 67 | Universal Open Pathogens Journal | Adyan Academic Press |
| 68 | US Open Clinical Pathology Journal | American Research Publications |
| 69 | World journal of pathology | Narain Publishers Pvt. Ltd (NPPL) |

**Table 3:** A list of Legitimate Journals in pathology (n = 89)

|  | Journal | Publisher |
| --- | --- | --- |
| 1 | Academic Pathology | Sage Publications |
| 2 | Acta Neuropathologica | Springer |
| 3 | Advances in Anatomic Pathology | Lippincott Williams & Wilkins |
| 4 | Alzheimer Disease & Associated Disorders | Lippincott Williams & Wilkins |
| 5 | The American Journal of Clinical Pathology | Oxford Academic |
| 6 | The American Journal of Forensic Medicine and Pathology | Lippincott Williams & Wilkins |
| 7 | The American Journal of Pathology | Elsevier |
| 8 | The American Journal of Surgical Pathology | Wolters Kluwer Health |
| 9 | Analytical Cellular Pathology | Hindawi Publishing Corporation |
| 10 | Annales de Pathologie | Elsevier |
| 11 | Annals of Diagnostic Pathology | Elsevier |
| 12 | Annual Review of Pathology-Mechanisms of Disease | Annual Reviews |
| 13 | APMIS | Wiley |
| 14 | Applied Immunohistochemistry & Molecular Morphology | Lippincott Williams & Wilkins |
| 15 | Archives of Pathology & Laboratory Medicine | College of American Pathologists |
| 16 | BMC Clinical Pathology | BioMed Central |
| 17 | Brain Pathology | Wiley |
| 18 | Brain Tumor Pathology | Springer |
| 19 | Cancer Cytopathology | Wiley |
| 20 | Cardiovascular Pathology | Elsevier |
| 21 | Case Reports in Pathology | Hindawi Publishing Corporation |
| 22 | Cellular Oncology | Springer |
| 23 | Clinical Medicine Insights: Pathology | Sage Publications |
| 24 | Clinical Neuropathology | Dustri-Verlag |
| 25 | CytoJournal | Medknow Publications |
| 26 | Cytometry Part B: Clinical Cytometry | Wiley |
| 27 | Cytopathology | Wiley |
| 28 | Diagnostic Cytopathology | Wiley |
| 29 | Diagnostic Histopathology | Elsevier |
| 30 | Diagnostic Pathology | BioMed Central |
| 31 | Disease Markers | Hindawi Publishing Corporation |
| 32 | Disease Models & Mechanisms | Cambridge |
| 33 | Endocrine Pathology | Springer |
| 34 | Experimental and Molecular Pathology | Elsevier |
| 35 | Experimental and Toxicologic Pathology | Elsevier |
| 36 | Expert Review of Molecular Diagnostics | Taylor & Francis Online |
| 37 | Fetal and Pediatric Pathology | Taylor & Francis Online |
| 38 | Folia Neuropathologica | Termedia |
| 39 | Forensic Science Medicine and Pathology | Springer |
| 40 | Histology and Histopathology | Histology and Histopathology |
| 41 | Histopathology | Wiley |
| 42 | HLA | Wiley |
| 43 | Human Pathology | Elsevier |
| 44 | Human Pathology: Case Reports | Medknow Publications |
| 45 | Indian Journal of Pathology and Microbiology | Medknow Publications |
| 46 | International Journal of Experimental Pathology | Wiley |
| 47 | International Journal of Gynecological Pathology | Dan Pasquarello |
| 48 | International Journal of Immunopathology and Pharmacology | Biomedical Research Press |
| 49 | International Journal of Surgical Pathology | Sage Publications |
| 50 | ISRN Pathology | Hindawi Publishing Corporation |
| 51 | Journal of Clinical Pathology | BMJ Publishing Group |
| 52 | Journal of Comparative Pathology | Elsevier |
| 53 | Journal of Cutaneous Pathology | Wiley |
| 54 | Journal of Hematopathology | Springer |
| 55 | The Journal of Molecular Diagnostics | Elsevier |
| 56 | Journal of Neuropathology & Experimental Neurology | Oxford University Press |
| 57 | Journal of Oral Pathology & Medicine | Wiley |
| 58 | The Journal of Pathology | Wiley |
| 59 | Journal of Pathology and Translational Medicine | The Korean Society of Pathologists |
| 60 | Journal of Pathology Informatics | Medknow Publications |
| 61 | Journal of Toxicologic Pathology | Japanese Society of Toxicologic Pathology |
| 62 | Laboratory Investigation | Nature |
| 63 | Leprosy Review | Lepra |
| 64 | Malaysian Journal of Pathology | Academy of Medicine of Malyasia |
| 65 | Medical Molecular Morphology | Springer |
| 66 | Modern Pathology | Nature |
| 67 | Neuropathology | Wiley |
| 68 | Neuropathology and Applied Neurobiology | Wiley |
| 69 | Pathobiology | Karger Publishers |
| 70 | Der Pathologe | Springer |
| 71 | Pathologie Biologie | Elsevier |
| 72 | Pathology | Elsevier |
| 73 | Pathology & Oncology Research | Springer |
| 74 | Pathology and Laboratory Medicine International | Dove Press |
| 75 | Pathology International | Wiley |
| 76 | Pathology Research and Practice | Elsevier |
| 77 | Pathology Research International | Hindawi Publishing Corporation |
| 78 | Patologiâ | Elsevier |
| 79 | Pediatric and Developmental Pathology | Sage Publications |
| 80 | Polish Journal of Pathology | Termedia |
| 81 | Science & Justice | Elsevier |
| 82 | Seminars in Diagnostic Pathology | Elsevier |
| 83 | Seminars in Immunopathology | Springer |
| 84 | The Journal of Pathology: Clinical Research | Wiley |
| 85 | Tissue Antigens | Wiley |
| 86 | Toxicologic Pathology | Sage Publications |
| 87 | Ultrastructural Pathology | Taylor & Francis Online |
| 88 | Veterinary Pathology | Sage Publications |
| 89 | Virchows Archiv | Springer |

All potential predatory journals in pathology shared at least one common poor-quality characteristics: lack of web-site integrity (n = 21, 31%) vs. (n = 0% in OA legitimate), missing/pending ISSN number (n = 36, 52%) vs. (n = 0% in OA legitimate), unreal or small number of issues per year (n = 22, 32%) vs. (n = 3, 3% in OA legitimate), emphasis on OA policy (n = 40, 58%) vs. (n = 32, 38% in OA legitimate), missing editorial board (n = 20, 29%) vs. (n = 1, 1% in OA legitimate), and ambiguous or unclear peer-review process (n = 38, 55%) vs. (n = 12, 14% in OA legitimate). Moreover, the majority (77%) of potential predatory journals accepted manuscript submissions via email. Absence of retraction, plagiarism, and copyright policies were all characteristics of the suspected journals as shown in Table 4 and Figure 1.

**Table 4:** A comparison of key characteristics of potential predatory journals/publishers in pathology based on the criteria proposed by Shamseer et al. (2)

| Criteria |  | Potential predatory*<br>N = 69, n (%) | Legitimate*<br>N = 85, n (%) |
| --- | --- | --- | --- |
| Aims and scope | Scope of interest: |  |  |
|  | a) Biomedical | 65 (94) | 82 (96) |
|  | b) Biomedical & non-biomedical | 2 (3) | 3 (4) |
| Journal name and publisher | Title similar to legitimate journal | 21 (30) |  |
| Home page integrity | The website contains spelling and/or grammar errors | 19 (28) | 3 (4) |
|  | Distorted and/or fuzzy images | 17 (25) | 2 (2) |
|  | Type of users targeted by homepage language: |  |  |
|  | a) Author | 46 (67) | 67 (79) |
|  | b) General | 21 (30) | 18 (21) |
| Contact information | The contact email address is non-journal affiliated (e.g., @hotmail.com) | 9 (13) | 2 (2) |
| Indexing and impact factor | Index Copernicus value | 13 (19) | 3 (4) |
|  | PubMed indexed | 0 (0) | 83 (98) |
|  | Medline indexed | 0 (0) | 69 (81) |
|  | DOAJ indexed | 1 (1) | 19 (22) |
|  | ISSN identified | 33(48) | 85 (100) |
|  | WAME listed | 0 (0) | 1 (1) |
|  | COPE listed | 0 (0) | 63 (74) |
| Editorial processing and peer review | Named editor-in-chief | 34 (49) | 84 (99) |
|  | Named editorial board | 49 (71) | 84 (99) |
|  | Description of manuscript handling process | 27 (39) | 82 (96) |
|  | Submission system: |  |  |
|  | a) E-mail | 53 (77) | 0 (0) |
|  | b) Journal submission system | 45 (65) | 85 (100) |
|  | Rapid publication is promised | 12 (17) | 0 (0) |
|  | States using peer-review | 64 (93) | 85 (100) |
| Publication ethics and policies | Lack of retraction policy | 43 (62) | 18 (21) |
|  | Presence of plagiarism policy | 40 (58) | 79 (93) |
| Publication model and copyright | Digital preservation of content | 42 (61) | 85 (100) |
|  | APC amount: |  |  |
|  | a) Stated | 57 (83) | 83 (98) |
|  | b) Not found | 10 (14) | 2 (2) |
|  | Retain copyright of published articles: |  |  |
|  | a) Stated | 37 (54) | 77 (91) |
|  | b) Not found | 30 (43) | 8 (9) |

\*Open access journals

**Figure 1:**
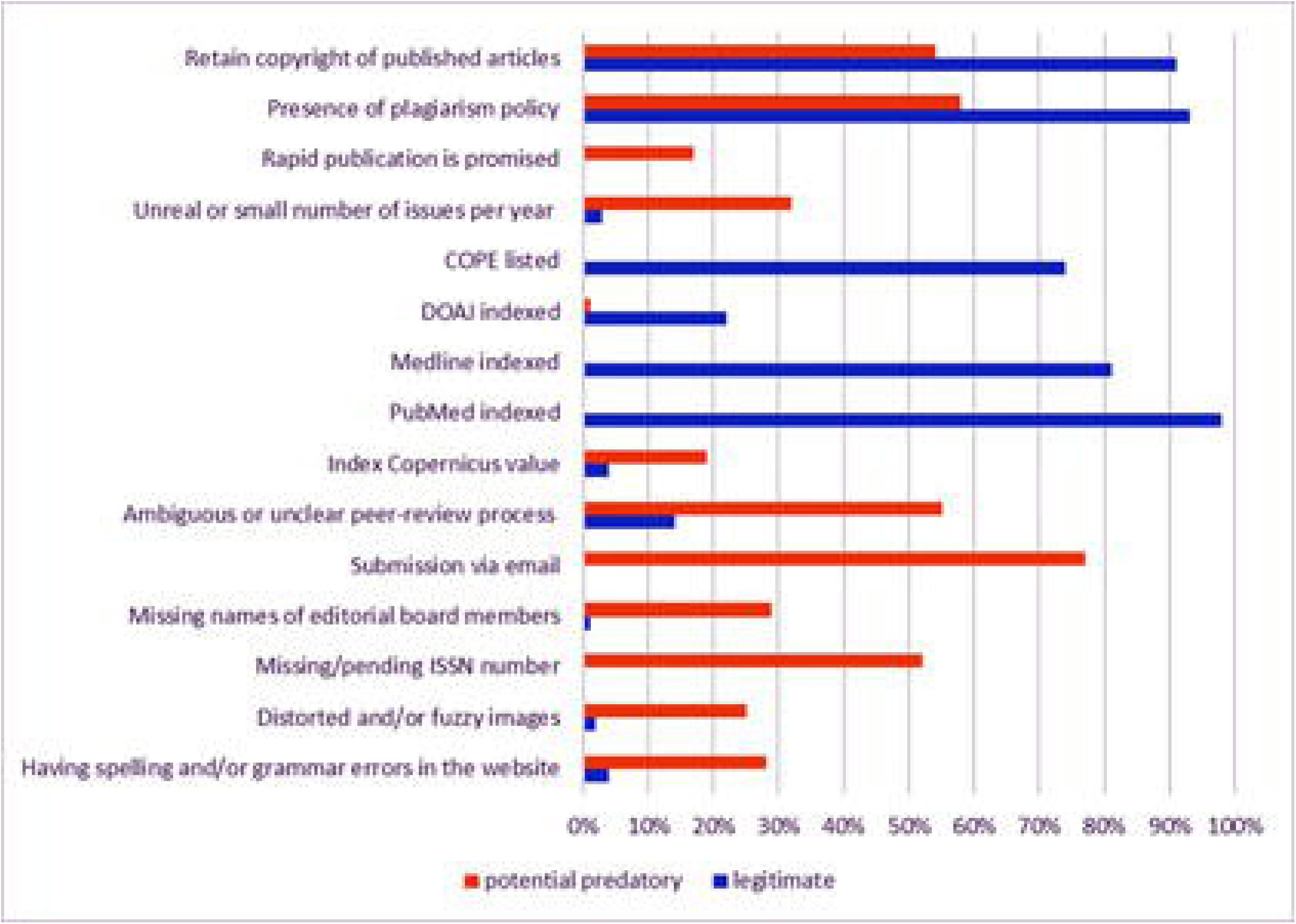
A comparison of quality characteristics among the pathology journals.

Furthermore, 21 (30%) potential predatory journals had misleading titles, which resemble or appear to be tied to those of legitimate ones (see Table 5). In addition, 31% of the potential predatory journals were indexed in the databases that generate bogus impact factors (e.g. Index Copernicus, Cosmos Impact Factor, and J-Gate). More specifically, 19% of the potential predatory journals presented their Index Copernicus value, whereas only 4% of the legitimate journals presented this impact factor.

**Table 5:**
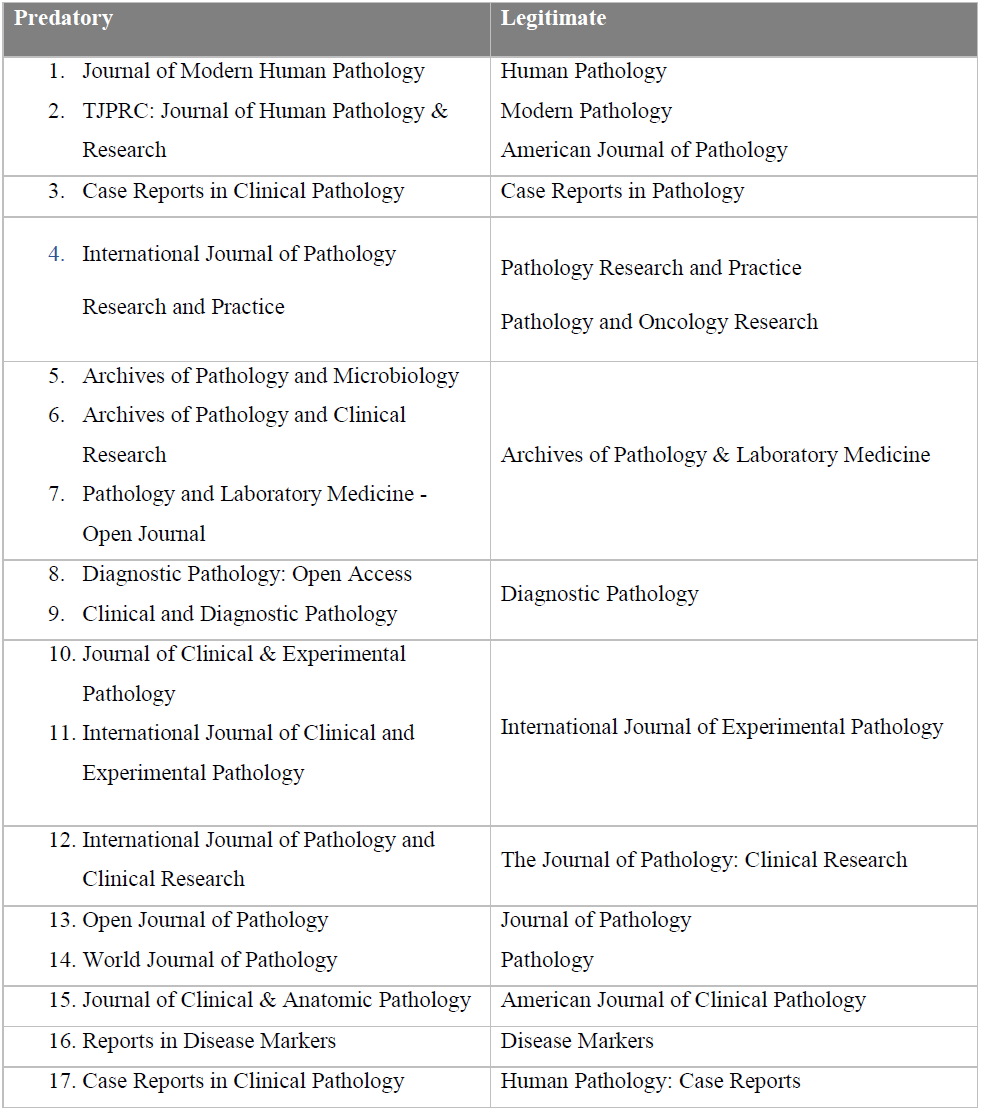
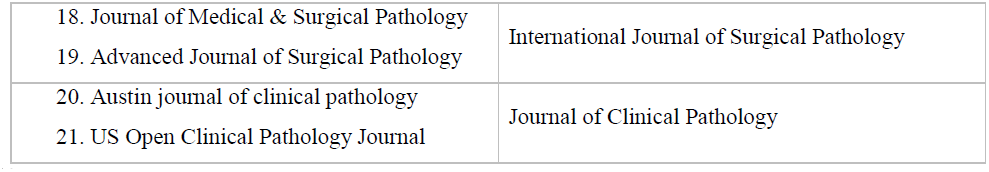
Potential predatory journals in pathology (n = 21) with names resembling/overlapping with those of the legitimate pathology journals

Eighty-three percent of the potential predatory journals displayed the required APCs on their web sites. The mean APC was significantly higher among the legitimate OA pathology journals in comparison with the predatory ones (US$ 2837.6 vs. US$ 814.3; range US$ 550-4100 vs. US$ 50-2700; p < 0.001).

Out of the suspected journals with valid ISSN (33/69), the vast majority (n = 23, 70%) of the targeted journals were originated from the United States, followed by India (n = 4, 12%). The remaining journals were distributed among the United Kingdom (n = 3, 10%), Nigeria (n = 2, 6%), and the United Arab Emirates (n = 1, 3%).

## Discussion

The Internet has dramatically transformed academic publishing, most notably, due to the introduction of OA publishing (2-5). Recently, there has been a rapid rise of online journals described as ‘predatory’ (3, 4). Such journals actively solicit manuscripts and charge publication fees without providing a robust peer review and proper editorial services (e.g. copyediting and proofreading) (2-4, 8, 9). Yet, it is important to note that OA is not correlated with the legitimacy of the journal.

In the present study, the impact of potential predatory journals in pathology was explored. Similar to other biomedical fields, our data indicate a substantial burden of such journals in academic pathology. One of the important findings of our study is that none of the potential predatory journals in pathology has been indexed in the major bibliographic databases such as PubMed/MEDLINE and Web of Science nor listed in COPE. These results are in line with a recent study on such journals in pediatrics (9). However, the studies in other biomedical fields (neurology/neurosurgery, physical medicine and emergency medicine) revealed the substantial contamination (up to 25%) with such journals in the major bibliographic databases (7, 12, 13).

Despite having none of the potential predatory journals indexed in PubMed, it is not a predatory-free database (14, 15). PubMed indexing policies are less strict compared with MEDLINE (14). In fact, 98% of the legitimate journals in this study were indexed in PubMed, whereas 81% were in MEDLINE. This might explain how some predatory journals managed to leak into the PubMed database (14). Although MEDLINE and PubMed are distinctively different databases, using the PubMed search engine queries both databases simultaneously (as well as PubMed Central).

Our study indicates that a substantial proportion (30%) of potential predatory journals in pathology may have similar names to the legitimate and renowned pathology journals (Table 5). In addition, predatory publishers usually send spam emails through which they invite authors to contribute to their journals and conferences promising a fast-track review and publishing (8). Summarized in Table 4 are the key characteristics of predatory journals/publishers (8), which may also guide the pathology researchers/scientists before deciding to submit an article to not well-known/unknown pathology journals (16).

Another strikingly distinct feature of potential predatory journals is APC (17). In line with previous studies in other medical disciplines, this study confirms that the mean of APC of potential predatory journals in pathology was significantly lower than that of legitimate OA journals in pathology (∼US$2837 vs. ∼US$814) (17). This marked difference along with “easy to publish” spam announcements might be a reason to attract some inexperienced and/or young academic pathologists to submit a paper to such journals.

Although the US and India were the countries of origin for the majority of the potential predatory journals, the methodology used in this paper provides better precision regarding the journal’s country of origin. In previous reports, Google Maps and 3D Street View were used predominantly to determine the journals and/or publishers’ country of origin based on the addresses displayed in their websites (7, 17). Such methodology is not credible to assess the country of origin, simply, because any random address can be presented in their web sites.

Notwithstanding its implications, two limitations to this study are important to highlight. First, the use of Beall’s list has been controversial as it was discontinued in 2017 and is consequently considered outdated (18, 19). Nevertheless, that list served as a start point given that each journal in our study was objectively apprised following the recommended criteria proposed by other researchers (8). Second, the Beall’s list was criticized for being subjective as it was established and updated by a single person; however, several well-conducted studies have relied on the list as an accessible reference for predatory journals (2, 6, 7, 17, 18). Finally, it is worth mentioning that the removal of Beall’s list led to the rise of new alternatives such as Cabell’s Blacklist (20).

In conclusion, this study highlights a substantial burden of potential predatory journals in pathology. A significant proportion of such journals (30%) have name of resemblance to the legitimate pathology journals, which may pose another significant challenge and threat to the academic community within this medical discipline. This study may aid pathology researchers in their decision-making process when submitting manuscripts for publication. Based on the obtained data, authors should check the journal’s status on PubMed/MEDLINE, Web of Science and DOAJ, as well as the previously proposed criteria, confirmed in this study, before submitting a manuscript to a pathology journal.

## Acknowledgment

This study was supported by the student grant number (#QUST-1-CMED-2018-10) provided by the College of Medicine, Qatar University. The preliminary data from the study were presented at the XXXII Congress of the International Academy of Pathology (IAP), October 2018, Amman, Jordan.

